## Supplementary material for "Metabolic pathway inference using multi-label classification with rich pathway features": S1 Appendix.

January 31, 2020

Here, we present analytical expression of mlLGPR model in Section 1.

### 1 Mathematical Derivations

We outline the process of deriving the objective cost function. The target function in our proposed model is the logistic regression, which represents the conditional probabilities through a non-linear logistic function $f(.)$ defined as:

$$f(\theta_j, \Phi(\mathbf{x}_i)) = p(\mathbf{y}_{i,j} = 1|\Phi(\mathbf{x}_i); \theta_j) = \frac{exp(\theta_j^\top \Phi(\mathbf{x}_i))}{exp(\theta_j^\top \Phi(\mathbf{x}_i)) + 1} \tag{1}$$

where $\mathbf{y}_{i,j}$ is the $j$-th element of the label vector $\mathbf{y}_i$ and $\theta_j$ is a $m$-dimensional weight vector for the $j$-th pathway that describes the space of $f(.)$ mapping from $\mathbb{R}^m$ to $2^{\mathcal{Y}}$ (a set of pathway space with $t$ possible pathways). The $\Phi(\mathbf{x}_i)$ is the (collective) transformation function that maps an instance $\mathbf{x}_i$ from $\mathbb{R}^d$ to a $\mathbb{R}^m$ dimensional vector. The Eq 1 can be written in more a compact form as:

$$p(\mathbf{y}_{i,j}|\Phi(\mathbf{x}_i); \theta_j) = f(\theta_j, \Phi(\mathbf{x}_i))^{\mathbf{y}_{i,j}}(1 - f(\theta_j, \Phi(\mathbf{x}_i)))^{1-\mathbf{y}_{i,j}} \tag{2}$$

Given $n$ training samples, the average likelihood of the $j$-th space parameters can be written as:

$$\begin{aligned} l(\theta_j) &= p(\mathbf{Y}|\Phi(\mathbf{X}); \theta_j) \\ &= \frac{1}{n}\prod_{i=1}^{n} p(\mathbf{y}_{i,j}|\Phi(\mathbf{x}_i); \theta_j) \\ &= \frac{1}{n}\prod_{i=1}^{n} (f(\theta_j, \Phi(\mathbf{x}_i))^{\mathbf{y}_{i,j}})(1 - f(\theta_j, \Phi(\mathbf{x}_i))^{1-\mathbf{y}_{i,j}}) \end{aligned} \tag{3}$$

where $\Phi(\mathbf{X}) = [\Phi(\mathbf{x}_1), \Phi(\mathbf{x}_2), ..., \Phi(\mathbf{x}_n)]^\top \in \mathbb{R}^{n \times m}$ represents the design matrix and $\mathbf{Y} = [\mathbf{y}_1, \mathbf{y}_2, ...\mathbf{y}_n]^\top$ is the label matrix. Taking the log-likelihood of the Eq 3 results in the following cost function:

$$ll(\theta_j) = \frac{1}{n}\sum_{i=1}^{n}(\mathbf{y}_{i,j}\theta_j^\top \Phi(\mathbf{x}_i) - \log(1 + \exp(\theta_j^\top \Phi(\mathbf{x}_i)))) \tag{4}$$

For $t$ pathways, we define the models weights matrix $\Theta = [\theta_1, \theta_2, ..., \theta_t]$ of size $m \times t$, and is estimated by maximizing the cost function from datasets. Thus, the Eq 4 can be generalized to $t$ pathways as:

$$C(\Theta) = \max_{\Theta} ll(\Theta) \tag{5}$$

Because jointly estimating $\Theta$ of Eq 5 in a straightforward way is intractable, we rather solve each weight vector $\theta_j$ of $\Theta$, individually. This optimization process is referred to as a *local optimization* technique:

$$C(\theta_j) = \max_{\theta_j} ll(\theta_j) \tag{6}$$

After adding a regularization term $\Omega(\theta_j)$ into Eq 6 and dropping the maximized term for notations brevity, the following objective function:

$$C(\theta_j) = ll(\theta_j) - \lambda\Omega(\theta_j) \tag{7}$$

The $\Omega(\theta_j)$ penalty term is an elastic-net, which is composed of two regularizers, namely $L_1$ and $L_2$, and plugging these two terms into Eq 7 we obtain:

$$C(\theta_j) = ll(\theta_j) - \lambda(\frac{1-\alpha}{2}||\theta_j||_2^2 + \alpha||\theta_j||_1) \tag{8}$$

where $||\theta_j||_2^2$ represents the $L_2$ regularizer while the term $||\theta_j||_1$ is $L_1$ regularizer. Both terms are controlled by $\alpha$. The $\lambda > 0$ is a hyper-parameter that controls the trade-off between $ll(\theta_j)$ and the two regularization terms.

Now, we want to choose $\theta_j$ so as to maximize $C(\theta_j)$. There are many optimizers to solve the cost function including coordinate descent algorithm (CD) [1]. However, we adopt mini-batch gradient descent (GD) (ascent in our adopted definition) ([2]) that converges much faster than CD [3]. The GD starts with some initial random guess for $\theta_j$, and repeatedly performs update to maximize the cost function $C(\theta_j)$, until the algorithm converges or reaches to a cutoff threshold:

$$\theta_j^{u+1} = \theta_j^u + \eta(\frac{\partial}{\partial\theta_j}C(\theta_j)) \tag{9}$$

Here, $\eta$ is called the learning rate and $u$ represents the current step. The update is simultaneously performed for all values of $j$, i.e., $(\theta_{j,1}, \theta_{j,2}, ..., \theta_{j,m})$. This is a very natural algorithm that repeatedly takes a step in the direction of steepest maximizing $C(\theta_j)$. To update the learning parameters, we take the partial derivatives on the right-hand side of the above equation. However, the rightmost term in Eq 8 is non-differentiable, making the equation non-smooth. As such, we resort to solve the Eq 8 with the $L_1$ and $L_2$ penalties, separately. The first two terms in the right part of Eq 8 are convex and differentiable. Let us denote the right part of Eq 8 as:

$$C_s(\theta_j) = ll(\theta_j) - \lambda\frac{1-\alpha}{2}||\theta_j||_2^2 \tag{10}$$

Taking the first-order derivative with respect to $\theta_j$ for Eq 10, we obtain the following formula:

$$\begin{aligned}
\frac{\partial}{\partial\theta_j}E_s(\theta_j) &= \frac{\partial}{\partial\theta_j}(ll(\theta_j) - \lambda\frac{1-\alpha}{2}||\theta_j||_2^2) \\
&= \frac{\partial}{\partial\theta_j}(\frac{1}{n}\sum_{i=1}^{n}(\mathbf{y}_{i,j}\theta_j^\top\Phi(\mathbf{x}_i) - \log(1+\exp(\theta_j^\top\Phi(\mathbf{x}_i)))) \\
&\quad - \lambda(1-\alpha)\theta_j \\
&= \frac{1}{n}\sum_{i=1}^{n}(\frac{\partial}{\partial\theta_j}(\mathbf{y}_{i,j}\theta_j^\top\Phi(\mathbf{x}_i) - \log(1+\exp(\theta_j^\top\Phi(\mathbf{x}_i)))) \\
&\quad - \lambda(1-\alpha)\theta_j \\
&= \frac{1}{n}\sum_{i=1}^{n}\Phi(\mathbf{x}_i)(\mathbf{y}_{i,j} - f(\theta_j, \Phi(\mathbf{x}_i))) - \lambda(1-\alpha)\theta_j
\end{aligned} \tag{11}$$

Because $\theta_{j,k} \neq 0$ in the rightmost term of Eq 8, we use the following defintion to obtain the gradient [4]:

$$\frac{\partial(-\lambda\alpha||\theta_j||_1)}{\partial\theta_j} = \begin{cases} \lambda\alpha & \text{if } \theta_{j,k} > 0 \\ -\lambda\alpha & \text{if } \theta_{j,k} < 0 \end{cases} \tag{12}$$

Denoting $\text{sign}(\theta_j)$ as the sign of the parameter vector $\theta_j$, the first order derivative of the rightmost in Eq 12 can be revised as:

$$\frac{\partial(-\lambda\alpha||\theta_j||_1)}{\partial\theta_j} = -\lambda\alpha \ \text{sign}(\theta_j) \tag{13}$$

Replacing Eq 11 and 13 in Eq 8, the first order derivatives of the objective cost function with respect to $\theta_j$ is:

$$\frac{\partial}{\partial\theta_j}C(\theta_j) = \frac{1}{n}\sum_{i=1}^{n}\Phi(\mathbf{x}_i)[\mathbf{y}_{i,j} - f(\theta_j, \Phi(\mathbf{x}_i))] - \lambda[(1-\alpha)\theta_j + \alpha \ \text{sign}(\theta_j)] \tag{14}$$

Putting Eq 14 into Eq 9, we obtain the final update algorithm:

$$\theta_j^{u+1} = \theta_j^u + \eta(\frac{1}{n}\sum_{i=1}^{n}\Phi(\mathbf{x}_i)[\mathbf{y}_{i,j} - f(\theta_j, \Phi(\mathbf{x}_i))] - \lambda[(1-\alpha)\theta_j + \alpha \ \text{sign}(\theta_j)]) \tag{15}$$
