## Supplementary material for "Metabolic pathway inference using multi-label classification with rich pathway features": S2 Appendix.

January 31, 2020

Given a set of ECs with abundance information, we define the following four sets of features: i)- reactions evidence features, ii)- pathways evidence features, iii)- pathway common features, and iv)- possible pathways features. Many features were re-designed from the work of Dale et. al [1], and were extracted according to the information available in MetaCyc database [2].

#### 1 Reactions Evidence Features

These real-valued features capture various reactions properties that are acquired from samples.

1. **fraction-total-ecs-to-distinct-ecs (numeric)**.
   The fraction of the total number of ECs, present in a sample, to the distinct set of ECs observed in that sample.

2. **fraction-total-possible-pathways-to-distinct-pathways (numeric)**
   The fraction of the total number of possible pathways that could be present in a sample to the distinct set of pathways represented in the total possible pathways.

3. **fraction-total-ecs-to-ecs-mapped-to-single-pathways (numeric)**
   The fraction of the total number of ECs that are associated with single pathways, to the total number of ECs, present in a sample.

4. **fraction-total-ecs-mapped-to-pathways (numeric)**
   The fraction of the total number of ECs, present in a sample, to the total number of pathways that are mapped according to MetaCyc database.

5. **fraction-total-distinct-ecs-contribute-in-subpathway-as-inside-superpathways (numeric)**
   The fraction of the total number of distinct ECs, contributing in pathways that are subpathways to superpathways according to MetaCyc database, to the total number of ECs, present in a sample.

6. **fraction-total-ecs-contribute-in-subpathway-as-inside-superpathways (numeric)**
   The fraction of the total number of ECs, contributing in pathways that are subpathways to superpathways according to MetaCyc database, to the total number of ECs, present in a sample.

7. **fraction-total-distinct-ecs-act-as-initial-reactions (numeric)**
   The fraction of the total number of distinct ECs, contributing at the beginning of pathways, to the total number of ECs, present in a sample.

8. **fraction-total-ecs-act-as-initial-reactions (numeric)**
   The fraction of the total number of ECs, contributing at the beginning of pathways, to the total number of ECs, present in a sample.

9. **fraction-total-distinct-ecs-act-as-final-reactions (numeric)**
   The fraction of the total number of distinct ECs, contributing at the end of pathways, to the total number of ECs, present in a sample.

10. **fraction-total-ecs-act-as-final-reactions (numeric)**
    The fraction of the total number of ECs, contributing at the end of pathways, to the total number of ECs, present in a sample.

11. **fraction-total-distinct-ecs-act-as-initial-and-final-reactions (numeric)**
    The fraction of the total number of distinct ECs, contributing at either the beginning or the end of pathways, to the total number of ECs, present in a sample.

12. **fraction-total-ecs-act-as-initial-and-final-reactions (numeric)**
The fraction of the total number of ECs, contributing at either the beginning or the end of pathways, to the total number of ECs, present in a sample.

13. **fraction-total-distinct-ecs-act-in-deg-or-detox-pathway (numeric)**
The fraction of the total number of distinct ECs that act either in degradation or detoxification pathways to the total number of ECs, present in a sample.

14. **fraction-total-ecs-act-in-deg-or-detox-pathway (numeric)**
The fraction of the total number of ECs that act either in degradation or detoxification pathways to the total number of ECs, present in a sample.

15. **fraction-total-distinct-ec-act-in-biosynthesis-pathway (numeric)**
The fraction of the total number of distinct ECs that act in biosynthetic pathways to the total number of ECs, present in a sample.

16. **fraction-total-ec-act-in-biosynthesis-pathway (numeric)**
The fraction of the total number of ECs that act in biosynthetic pathways to the total number of ECs, present in a sample.

17. **fraction-total-distinct-ec-act-in-energy-pathway (numeric)**
The fraction of the total number of distinct ECs that act in energy pathways to the total number of ECs, present in a sample.

18. **fraction-total-ec-act-in-energy-pathway (numeric)**
The fraction of the total number of ECs that act in energy pathways to the total number of ECs in a sample, present in a sample.

19. **fraction-total-ecs-to-total-reactions (numeric)**
The fraction of the total number of ECs to the total number of reactions that are catalyzed by enzymes, encoded as ECs, in a given sample.

20. **fraction-total-distinct-ecs-to-total-distinct-reactions (numeric)**
The fraction of the total number of distinct ECs to the total number of distinct reactions that are catalyzed by enzymes, encoded as ECs, in a given sample.

21. **fraction-total-ec-contribute-in-unique-reaction (numeric)**
The fraction of the total number of ECs to the total number of reactions unique to ECs that are catalyzed by enzymes in a given sample.

22. **fraction-total-distinct-ec-contribute-to-reactions-has-taxonomic-range (numeric)**
The fraction of the total number of distinct ECs that have taxonomic information to the total number of ECs, present in a sample.

23. **fraction-total-pathways-over-total-ecs (numeric)**
The fraction of the total number of possible pathways that could be present in a sample to the total number of ECs in that sample.

24. **fraction-total-pathways-over-distinct-ec (numeric)**
The fraction of the total number of possible pathways that could be present in a sample to the total number of distinct ECs in that sample.

25. **fraction-total-distinct-pathways-over-distinct-ec (numeric)**
The fraction of the total number of distinct possible pathways that could be present in a sample to the total number of distinct ECs in that sample.

26. **fraction-distinct-ec-contributes-in-subpathway-over-distinct-pathways (numeric)**
The fraction of the total number of distinct ECs, contributing in subpathways, to the total number of distinct possible pathways that could be present in a given sample.

27. **fraction-ec-contributes-in-subpathway-over-total-pathways (numeric)**
The fraction of the total number of ECs, contributing in subpathways, to the total number of possible pathways that could be present in a given sample.

28. **fraction-distinct-ec-act-in-deg-or-detox-pathway-over-distinct-pathways (numeric)**
The fraction of the total number of distinct ECs, acting in degradation or detoxification pathways, to the total number of possible distinct pathways that could be present in a given sample.

29. **fraction-distinct-ec-act-in-deg-or-detox-pathway-over-total-pathways (numeric)**
The fraction of the total number of distinct ECs, acting in degradation or detoxification pathways, to the total number of possible pathways that could be present in a given sample.

30. **fraction-ec-act-in-deg-or-detox-pathway-over-total-pathways (numeric)**
The fraction of the total number of ECs, acting in degradation or detoxification pathways, to the total number of possible pathways that could be present in a given sample.

31. **fraction-distinct-ec-act-in-biosynthesis-pathway-over-distinct-pathways (numeric)**
The fraction of the total number of distinct ECs, acting in biosynthetic pathways, to the total number of possible distinct pathways that could be present in a given sample.

32. **fraction-distinct-ec-act-in-biosynthesis-pathway-over-total-pathways (numeric)**
The fraction of the total number of distinct ECs, acting in biosynthetic pathways, to the total number of possible pathways that could be present in a given sample.

33. **fraction-ec-act-in-biosynthesis-pathway-over-total-pathways (numeric)**
The fraction of the total number of ECs, acting in biosynthetic pathways, to the total number of possible pathways that could be present in a given sample.

34. **fraction-distinct-ec-act-in-energy-pathway-over-distinct-pathways (numeric)**
The fraction of the total number of distinct ECs, acting in energy pathways, to the total number of possible distinct pathways that could be present in a given sample.

35. **fraction-distinct-ec-act-in-energy-pathway-over-total-pathways (numeric)**
The fraction of the total number of distinct ECs, acting in energy pathways, to the total number of possible pathways that could be present in a given sample.

36. **fraction-ec-act-in-energy-pathway-over-total-pathways (numeric)**
The fraction of the total number of ECs, acting in energy pathways, to the total number of possible pathways that could be present in a given sample.

37. **fraction-total-reactions-over-total-pathways (numeric)**
The fraction of the total number of reactions, catalyzed by enzymes, encoded as ECs, to the total number of possible pathways that could be present in a given sample.

38. **fraction-total-reactions-over-distinct-pathways (numeric)**
The fraction of the total number of reactions, catalyzed by enzymes and encoded as ECs, to the total number of possible distinct pathways that could be present in a given sample.

39. **fraction-distinct-reaction-over-distinct-pathways (numeric)**
The fraction of the total number of distinct reactions, catalyzed by enzymes and encoded as ECs, to the total number of possible distinct pathways that could be present in a given sample.

40. **ecs-in-energy-pathways-mostly-missing (numeric)**
The total number of energy pathways that have more than half of their true ECs mapping are missing in a given sample.

41. **ecs-in-pathways-mostly-present (numeric)**
The total number of pathways that have more than half of their true ECs mapping are present while missing only one ECs in a given sample.

42. **all-initial-ecs-present-in-pathways (numeric)**
The total number of pathways that have at least two of their beginning ECs are present in a given sample.

43. **all-final-ecs-present-in-pathways (numeric)**
The total number of pathways that have at least two of their final ECs are present in a given sample.

44. **all-initial-and-final-ecs-present-in-pathways (numeric)**
The total number of pathways that have at least two of their beginning and final ECs are present in a given sample.

45. **all-initial-ecs-present-in-deg-or-detox-pathways (numeric)**
The total number of degradation or detoxification pathways that have at least two of their beginning ECs are present in a given sample.

46. **all-final-ecs-present-in-deg-or-detox-pathways (numeric)**
The total number of degradation or detoxification pathways that have at least two of their final ECs are present in a given sample.

47. **all-initial-ecs-present-in-biosynthesis-pathways (numeric)**
The total number of biosynthetic pathways that have at least two of their beginning ECs are present in a given sample.

48. **all-final-ecs-present-in-biosynthesis-pathways (numeric)**
The total number of biosynthetic pathways that have at least two of their final ECs are present in a given sample.

49. **most-ecs-absent-in-pathways (numeric)**
The total number of pathways that have only one of their true ECs mapping is present in a given sample.

50. **most-ecs-absent-not-distinct-in-pathways (numeric)**
The total number of pathways that have half of their true ECs mapping are not distinct to pathways and missing in a given sample.

51. **one-ec-present-but-in-minority-in-pathways (numeric)**
The total number of pathways that have only one of their true ECs mapping is present and is considered minority to pathways in a given sample.

52. **all-distinct-ec-present-in-pathways (numeric)**
The total number of pathways that have all of their true distinct ECs mapping are present in a given sample.

53. **all-ecs-present-in-pathways (numeric)**
The total number of pathways that have all of their true ECs mapping are present in a given sample.

54. **all-distinct-ec-present-or-orphaned-in-pathways (numeric)**
The total number of pathways that have all of their true distinct ECs mapping are present or are orphaned according to MetaCyc in a given sample.

55. **all-ec-present-or-orphaned-in-pathways (numeric)**
The total number of pathways that have all of their true ECs mapping are present or are orphaned according to MetaCyc in a given sample.

56. **majority-of-ecs-absent-in-pathways (numeric)**
The total number of pathways that have more than half of their true ECs mapping are missing in a given sample.

57. **majority-of-ecs-present-in-pathways (numeric)**
The total number of pathways that have more than half of their true ECs mapping are present in a given sample.

58. **majority-of-distinct-ecs-present-in-pathways (numeric)**
The total number of pathways that have more than half of their true distinct ECs mapping are present in a given sample.

59. **majority-of-reactions-present-distinct-in-pathways (numeric)**
The total number of pathways that have more than half of their true ECs mapping are present and distinct to pathways in a given sample.

60. **missing-at-most-one-ec-in-pathways (numeric)**
The total number of pathways that have only one of their true ECs mapping is absent in a given sample.

61. **has-distinct-ecs-present-in-pathways (numeric)**
The total number of pathways that some of their true distinct ECs mapping are present in a given sample.

62. **fraction-distinct-ecs-present-or-orphaned-in-pathways (numeric)**
The total fraction of distinct ECs or orphaned reactions associated to the possible pathways in a given sample to the pathways true ECs mapping.

63. **fraction-reactions-present-or-orphaned-distinct-in-pathways (numeric)**
The total fraction of ECs or orphaned reactions that are distinctly associated to possible pathways in a given sample to the pathways true ECs mapping.

64. **fraction-reactions-present-or-orphaned-in-pathways (numeric)**
The total fraction of ECs or orphaned reactions associated to the possible pathways in a given sample to the pathways true ECs mapping.

65. **num-distinct-reactions-present-or-orphaned-in-pathways (numeric)**
The total number of distinct ECs or orphaned reactions associated to the possible pathways are present in a given sample.

66. **num-reactions-present-or-orphaned-in-pathways (numeric)**
The total number of ECs or orphaned reactions associated to the possible pathways are present in a given sample.

67. **evidence-info-content-norm-present-in-pathways (numeric)**
The total evidence information content of pathways, normalized by the number of reactions associated with pathways that are present in a given sample. For a single pathway $y_j$ of sample $\mathbf{x}_i$, this feature is computed as:

$$
\begin{aligned}
evidence &= \frac{1}{\sum_{\hat{a} \in \mathbf{y}_j^{(i)}} \hat{a}} \sum_{a \in \mathbf{x}^{(i)}} \frac{1}{\sum_{y_k \in \mathcal{Y}} \sum_{(e,a') \in y_k} \delta(a')} \\
\text{where } \delta(a') &= \begin{cases} 1, & \text{if } a' \geq 1 \text{ and } a' \in \mathbb{N} \\ 0, & \text{otherwise} \end{cases}
\end{aligned} \tag{1}
$$

where $\mathcal{Y}$ represents the universal set of pathways while $e$ represents an EC and $a$, $a'$ and $\hat{a}$ are abundances in $\mathbf{x}^{(i)}$, $y_k$, and $\mathbf{y}_j^{(i)}$, respectively and are elements in $\mathbb{N}$. This feature measures how strongly the evidence for the pathway $j$ based on ECs. The ECs contributing to many pathways have low evidence for the presence of the reactions that EC catalyze.

68. **evidence-info-content-present-in-pathways (numeric)**
The total evidence information content of pathways in a given sample.

### 2 Pathways Evidence Features

These features are designed to capture simple patterns for each pathway from samples. They are combination of two types: boolean and numeric.

1. **ecs-mostly-present-in-pathway (boolean)**
True if a pathway is: a)- missing at most one EC and b)- half of its ECs is present.

2. **prob-ecs-mostly-present-in-pathway (numeric)**
The fraction of the total number of ECs associated to a pathway, present in a sample, to the true ECs mapping of that pathway, satisfying two conditions: a)- missing at most one EC and b)- half of that pathway's ECs is present.

3. **all-initial-ecs-present-in-pathway (boolean)**
True if the first two ECs associated to a pathway are present in a sample.

4. **prob-initial-ecs-present-in-pathway (numeric)**
The fraction of the first two ECs associated to a pathway, if present in a sample, to the first two true ECs mapping of that pathway.

5. **all-final-ecs-present-in-pathway (boolean)**
True if the last two ECs associated to a pathway are present in a sample.

6. **prob-final-ecs-present-in-pathway (numeric)**
The fraction of the last two ECs associated to a pathway, if present in a sample, to the last two true final ECs mapping of that pathway.

7. **all-initial-and-final-ecs-present-in-pathway (boolean)**
True if the first two and the last two ECs associated to a pathway are present in a sample.

8. **prob-all-initial-and-final-ecs-present-in-pathway (numeric)**
The fraction of the first two and the last two ECs associated to a pathway, if present in a sample, to the first two and the last two true ECs mapping of that pathway.

9. **all-initial-ecs-present-in-deg-or-detox-pathway (boolean)**
True if the first two ECs associated to a degradation or a detoxification pathway are present in a sample.

10. **prob-all-initial-ecs-present-in-deg-or-detox-pathway (numeric)**
The fraction of the first two ECs associated to a degradation or a detoxification pathway, if present in a sample, to the first two true ECs mapping of that pathway.

11. **all-initial-ecs-present-in-biosynthesis-pathway (boolean)**
True if the first two ECs associated to a biosynthetic pathway are present in a sample.

12. **prob-all-initial-ecs-present-in-biosynthesis-pathway (numeric)**
The fraction of the first two ECs associated to a biosynthetic pathway, if present in a sample, to the first two true ECs mapping of that pathway.

13. **most-ecs-absent-in-pathway (boolean)**
True if only one EC associated to a pathway is present in a sample.

14. **most-ecs-absent-not-distinct-in-pathway (boolean)**
True if half of ECs associated to a pathway in a sample is not distinct to that pathway.

15. **one-ec-present-but-in-minority-in-pathway (boolean)**
True if only one EC associated to a pathway is present in a sample and is considered a minority to that pathway.

16. **all-distinct-ec-present-in-pathway (boolean)**
True if all distinct ECs associated to a pathway are present in a sample.

17. **all-ecs-present-in-pathway (boolean)**
True if all ECs associated to a pathway are present in a sample.

18. **all-distinct-ec-present-or-orphaned-in-pathway (boolean)**
True if all distinct ECs associated to a pathway are present in a sample or orphaned according to MetaCyc.

19. **all-ec-present-or-orphaned-in-pathway (boolean)**
True if all ECs associated to a pathway are present in a sample or orphaned according to MetaCyc.

20. **majority-of-ecs-absent-in-pathway (boolean)**
True if more than half of ECs associated to a pathway in a sample are missing.

21. **majority-of-ecs-present-in-pathway (boolean)**
True if more than half of ECs associated to a pathway in a sample are present.

22. **majority-of-distinct-ecs-present-in-pathway (boolean)**
True if more than half of distinct ECs associated to a pathway in a sample are present.

23. **majority-of-reactions-present-distinct-in-pathway (boolean)**
True if more than half of ECs associated to a pathway in a sample are present and distinct to that pathway.

24. **missing-at-most-one-ec-in-pathway (boolean)**
True if only one EC associated to a pathway in a sample is missing.

25. **has-distinct-ecs-present-in-pathway (boolean)**
True if some distinct ECs associated to a pathway in a sample are present.

26. **fraction-distinct-ecs-present-or-orphaned-in-pathway (numeric)**
The fraction of distinct ECs or orphaned reactions associated to a pathway in a sample.

27. **fraction-reactions-present-or-orphaned-distinct-in-pathway (numeric)**
The fraction of ECs or orphaned reactions that are distinctly associated to a pathway in a sample.

28. **fraction-reactions-present-or-orphaned-in-pathway (numeric)**
The fraction of ECs or orphaned reactions associated to a pathway in a sample.

29. **num-distinct-reactions-present-or-orphaned-in-pathway (numeric)**
The number of distinct ECs or orphaned reactions associated to a pathway is present in a sample.

30. **num-reactions-present-or-orphaned-in-pathway (numeric)**
The number of ECs or orphaned reactions associated to a pathway is present in a sample.

31. **evidence-info-content-norm-present-in-pathway (numeric)**
    The total evidence information content of a pathway, normalized by the number of reactions associated with that pathways which are present in a sample.

32. **evidence-info-content-present-in-pathway (numeric)**
    The total evidence information content of a pathway in a sample.

### 3 Pathway Common Features

This feature set is designed to recognize (mis-)matches between a list of ECs from samples and the true mappings of pathways to ECs.

1. **ec-pathway-common-present (boolean)**

### 4 Possible Pathways Features

This feature set is of two types: i)- a boolean representation indicating the presence/absence of pathways in samples that exceed a user-defined cutoff threshold (0.5 in our setting), and ii)- a numerical representation encoding the probabilities of pathways to be present in samples.

1. **possible-pathways-present (boolean)**

2. **prob-possible-pathways-present (numeric)**
