## Supplementary material for "Metabolic pathway inference using multi-label classification with rich pathway features": S3 Appendix.

January 31, 2020

#### 1 Dataset Characteristics

Defining benchmark corpora for pathway prediction methods will promote thorough understanding of critical factors involved in the pathway predictions' performances. As such, we developed 12 benchmark datasets used in the experiments, with detailed characteristics summarized in Table 1. The 12 datasets cover a wide range of cases with diverse multi-label properties, ranging from synthetic to single organism to multiple organisms. Therefore, the experimental studies published in this paper are comprehensive, and aim to provide a strong basis for rigorous assessment of pathway prediction algorithms' effectiveness. For each dataset $\mathcal{S}$, we use $|\mathcal{S}|$ and $\text{L}(\mathcal{S})$ to represent the number of instances and pathway labels, respectively. In addition, we also present some characteristics of the multi-label datasets, which are denoted as:

1. Label cardinality ($\text{LCard}(\mathcal{S}) = \frac{1}{n}\sum_{i=1}^{i=n}\sum_{j=1}^{j=t}\mathbb{I}[\mathbf{Y}_{i,j} \neq -1]$), where $\mathbb{I}$ is an indicator function. It denotes the average number of pathways in $\mathcal{S}$.

2. Label density ($\text{LDen}(\mathcal{S}) = \frac{LCard(\mathcal{S})}{L(\mathcal{S})}$). This is simply obtained through normalizing $\text{LCard}(\mathcal{S})$ by the number of total pathways in $\mathcal{S}$.

3. Distinct label sets ($\text{DL}(\mathcal{S})$). This notation indicates the number of distinct pathways in $\mathcal{S}$.

4. Proportion of distinct label sets ($\text{PDL}(\mathcal{S}) = \frac{DL(\mathcal{S})}{|\mathcal{S}|}$). It represents the normalized version of $\text{DL}(\mathcal{S})$, and is obtained by dividing DL(.) with the number of instances in $\mathcal{S}$.

The notations $\text{R}(\mathcal{S})$, $\text{RCard}(\mathcal{S})$, $\text{RDen}(\mathcal{S})$, $\text{DR}(\mathcal{S})$, and $\text{PDR}(\mathcal{S})$ have similar meanings as before but for the enzymatic reactions $\mathcal{E}$ in $\mathcal{S}$, and $\text{PLR}(\mathcal{S})$ represent a ratio of $\text{L}(\mathcal{S})$ to $\text{R}(\mathcal{S})$. The experimental multi-label datasets can be compartmentalized into three groups based on the process of their curation, ordered by increasing purity, as: 1)-*golden* (Section 1.1), 2)- symbiotic data (Section 1.2), 3)-*CAMI* low complexity data (Section 1.3), 4)-HOT metagenomics dataset (Section 1.4), and 5)- *synthetic* datasets (Section 1.5).

##### 1.1 Golden Dataset

The golden dataset can be decomposed into two tiers. Tier 1 dataset composed from six databases, retrieved from biocyc website: *EcoCyc (v21)*, *HumanCyc (v19.5)*, *AraCyc (v18.5)*, *YeastCyc (v19.5)*, *LeishCyc (v19.5)*, and *TrypanoCyc (v18.5)*, and are refined to include only those pathways that cross-intersect with the *MetaCyc* database v21 [1]. For each of these databases, we extracted both the enzymatic reactions and the associated pathways to obtain the golden tier 1 data.

Tier 2 dataset, referred to as *SixDB*, is comprised of the previous six databases, by performing enumeration to create all possible combinations, which resulted in 63 samples, using the following formula:

$$|\mathcal{S}| = \sum_{k=1}^{k=G}\binom{G}{k} \tag{1}$$

where $|.|$ denotes the number of samples in $\mathcal{S}$ and $G$ is the number of databases, which is 6. While the biological contexts in this data were excluded, the pathways were retained. Therefore, it is not considered as pure as Tier 1 golden dataset.

To better resolve the intersected pathways among the six datasets, we used UpSet [2, 3] to encode all intersections of six datasets. Fig 1 summarizes the results where the columns of the matrix use binary circled-shaped patterns to define the applied intersected datasets, and the bars, just above the matrix columns, represent the number of elements in each intersection. The bars at the bottom left, plotted along the rows of the matrix, provide information regarding the total intersection size of a dataset. Also, Fig 1 shows several interesting observations, for example, LeishCyc has the lowest number in both: the distinct pathways, having only 4 pathways, and the aggregated number of pathways from all enumeration of intersected sets, which represents

| Dataset | $\lvert\mathcal{S}\rvert$ | L($\mathcal{S}$) | LCard($\mathcal{S}$) | LDen($\mathcal{S}$) | DL($\mathcal{S}$) | PDL($\mathcal{S}$) | R($\mathcal{S}$) | RCard($\mathcal{S}$) | RDen($\mathcal{S}$) | DR($\mathcal{S}$) | PDR($\mathcal{S}$) | PLR($\mathcal{S}$) | Domain |
|---|---|---|---|---|---|---|---|---|---|---|---|---|---|
| AraCyc | 1 | 510 | 510 | 1 | 510 | 510 | 2182 | 2182 | 1 | 1034 | 1034 | 0.2337 | Arabidopsis thaliana |
| EcoCyc | 1 | 307 | 307 | 1 | 307 | 307 | 1134 | 1134 | 1 | 719 | 719 | 0.2707 | Escherichia coli K-12 substr.MG1655 |
| HumanCyc | 1 | 279 | 279 | 1 | 279 | 279 | 1177 | 1177 | 1 | 693 | 693 | 0.2370 | Homo sapiens |
| LeishCyc | 1 | 87 | 87 | 1 | 87 | 87 | 363 | 363 | 1 | 292 | 292 | 0.2397 | Leishmania major Friedlin |
| TrypanoCyc | 1 | 175 | 175 | 1 | 175 | 175 | 743 | 743 | 1 | 512 | 512 | 0.2355 | Trypanosoma brucei |
| YeastCyc | 1 | 229 | 229 | 1 | 229 | 229 | 966 | 966 | 1 | 544 | 544 | 0.2371 | Saccharomyces cerevisiae |
| Symbiotic | 3 | 119 | 39.6667 | 0.3333 | 59 | 19.6667 | 304 | 101.3333 | 0.3333 | 130 | 43.3333 | 0.3914 | Composed of Moranella and Tremblaya |
| CAMI | 40 | 6261 | 156.5250 | 0.0250 | 674 | 16.8500 | 14269 | 356.7250 | 0.0250 | 1083 | 27.0750 | 0.4388 | Simulated microbiomes of low complexity |
| HOT | 4 | 2178 | 311.1429 | 0.1429 | 781 | 111.5714 | 182675 | 26096.4286 | 0.1429 | 1442 | 206.0000 | 0.0119 | Metagenomic Hawaii Ocean Time-series (10m, 75m, 110m, and 500m) |
| Synset-1 | 15000 | 6801364 | 453.4243 | 0.00007 | 2526 | 0.1684 | 30901554 | 2060.1036 | 0.00007 | 3650 | 0.2433 | 0.2201 | Synthetically generated (uncorrupted) |
| Synset-2 | 15000 | 6806262 | 453.7508 | 0.00007 | 2526 | 0.1684 | 34006386 | 2267.0924 | 0.00007 | 3650 | 0.2433 | 0.2001 | Synthetically generated (corrupted) |

Table 1: **Characteristics of the experimental datasets.** The notations $\lvert\mathcal{S}\rvert$, L($\mathcal{S}$), LCard($\mathcal{S}$), LDen($\mathcal{S}$), DL($\mathcal{S}$), and PDL($\mathcal{S}$) represent number of instances, number of pathway labels, pathway labels cardinality, pathway labels density, distinct pathway labels set, and proportion of distinct pathway labels set for $\mathcal{S}$, respectively. The notations R($\mathcal{S}$), RCard($\mathcal{S}$), RDen($\mathcal{S}$), DR($\mathcal{S}$), and PDR($\mathcal{S}$) have similar meanings as before but for the enzymatic reactions $\mathcal{E}$ in $\mathcal{S}$. PLR($\mathcal{S}$) represents a ratio of L($\mathcal{S}$) to R($\mathcal{S}$). The last column denotes the domain of $\mathcal{S}$.

the cardinality of LeishCyc pathways, i.e., 87 pathways (see Table 1), while AraCyc data has the highest number in both categories (271 distinct pathways and 510 the aggregated number of pathways). These observations were substantially beneficial during our experimental inspections.

#### 1.2 Symbiotic Dataset

The symbiotic data illustrates the dual bacterial symbionts of mealybug *Planococcus citri* composed of *Candidatus Moranella endobia* (GenBank NC-015735) living inside *Candidatus Tremblaya princeps* (GenBank NC-015736) [4]. We used MetaPathways v2.5 ([5]) and Pathway Tools version 21 to generate ePGDBs with the default settings. The symbiotic *Candidatus Moranella endobia* and *Candidatus Tremblaya princeps* genomes can be downloaded from GenBank under accession numbers NC-015735 and NC-015736).

#### 1.3 CAMI Dataset

The CAMI (Critical Assessment of Metagenome Interpretation) low complexity dataset [6] is a simulated dataset from 40 genomes of low complexity. The dataset has various purposes related to evaluating the performances of assembly, profiling, and binning tools. This dataset is placed in a lower order of purity than the previous golden samples because it constitutes synthetic mock community of microbiomes. Similar to symbiotic data, MetaPathways v2.5 ([5]) is employed to generate ePGDBs. The simulated CAMI low complexity dataset can be obtained from edwards.sdsu.edu/research/cami-challenge-datasets/.

Figure 1: **Matrix layout for all possible intersections among EcoCyc, HumanCyc, AraCyc, YeastCyc, LeishCyc, and TrypanoCyc dataset.** Brown circles in the matrix indicate sets that are part of the intersection and their distributions are shown as a vertical bar above the matrix while the aggregated number of pathways from intersected sets for each sample is represented by a horizontal bar at the bottom left. More information is provided in Table 1.

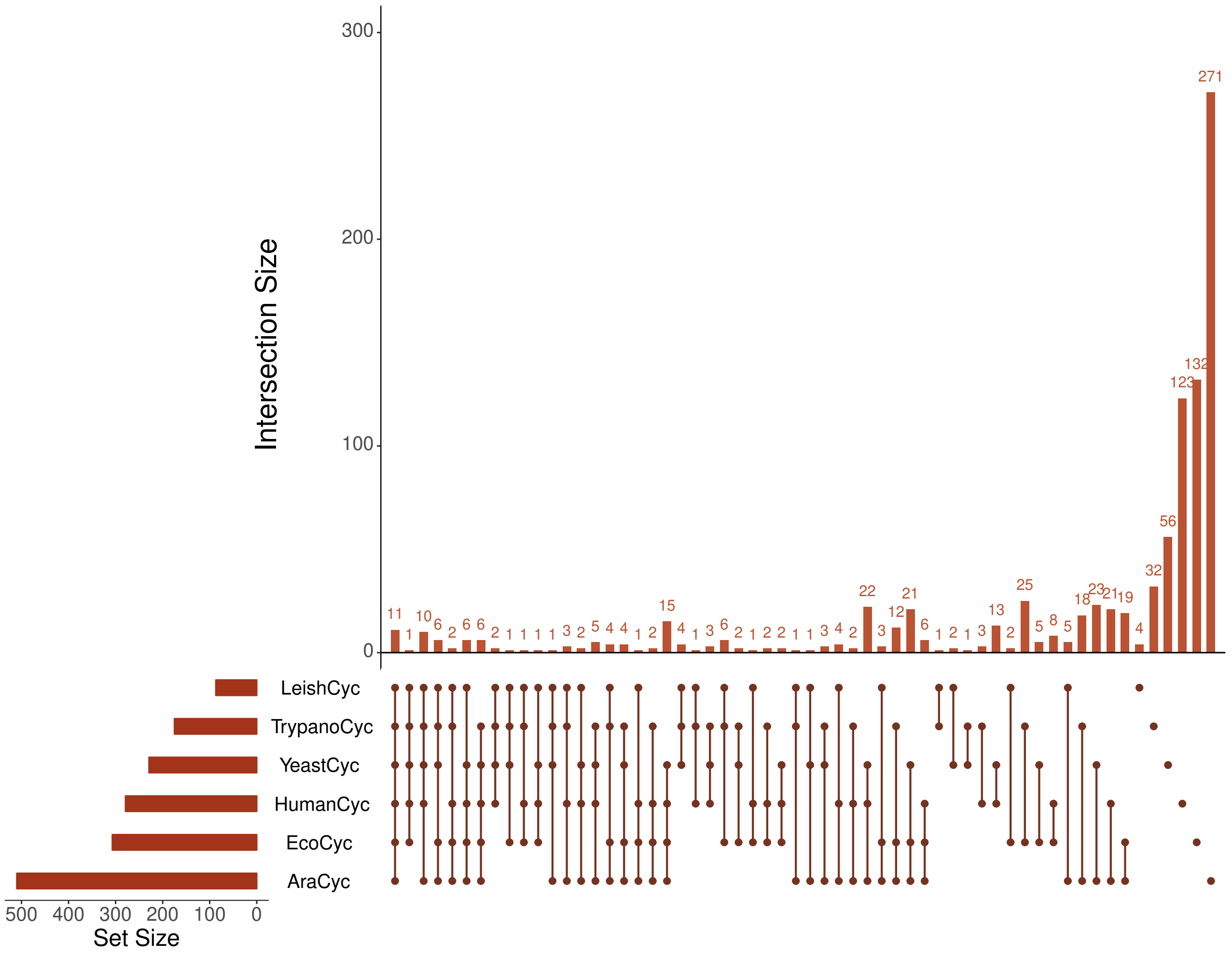


#### 1.4 HOT Dataset

The HOT metagenome (DNA) dataset is composed of complex microbial communities from 25m, 75m, 110m (sunlit) and 500m (dark) ocean depth intervals [7]. Unassembled metagenomic pyrosequences from the Hawaii Ocean Time-series (10m, 75m, 110m, and 500m) can be obtained from the NCBI Sequence Read Archive under accession numbers SRX007372, SRX007369, SRX007370, SRX007371. To generate ePGDBs, MetaPathways v2.5 ([5]) is employed.

#### 1.5 Synthetic Samples Generation

The *in silico* synthetic dataset is constructed by selecting a list of pathways, at first, then creating instances to curate a dataset. This dataset is used to train and evaluate the mlLGPR's predictive performance. The data generation process can be summarized in three key stages:

- **Phase 1: Specifying Pathways.** We load all the available pathways from the MetaCyc database and tier 1 data. Then, we keep a list of pre-specified pathways to be examined while truncating the rest. The considered pathway list $\mathcal{Y}$ is used for training and performance evaluation.

- **Phase 2: Generation Process.** We construct an instance by randomly selecting a subset of pathways from $\mathcal{Y}$, i.e., $\widehat{\mathcal{Y}}_i \subset \mathcal{Y}$. Given $\widehat{\mathcal{Y}}_i$, we perform mapping onto MetaCyc to retrieve a list of enzymatic reactions with abundances so as to generate an instance $\mathbf{x}^{(i)}$. Together $(\mathbf{x}^{(i)}, \mathbf{y}^{(i)})$ forms a synthetic sample. Replicating this process $n$ times results in a dataset $\mathcal{S} = \{(\mathbf{x}^{(i)}, \mathbf{y}^{(i)})|1 < i \leq n\}$. The enzymatic reactions are indicated by the EC (Enzyme Commission) numbers, which denote the numerical classification of enzymes based on the reactions they catalyze. In the experiment, we consider all EC numbers, including the incomplete ones, such as EC 1.2.3.-.

- **Phase 3: Corruption Process.** The corruption is explicitly applied by first selecting a sample $(\mathbf{x}^{(i)}, \mathbf{y}^{(i)})$, uniformly, from a newly created $\mathcal{S}$. Then, for each pathway $y_j \in \mathbf{y}^{(i)}$, one of the three options is selected: i)- retain $y_j$, ii)- remove a list of enzymatic reactions associated with $y_j$, or iii)- insert a list of false enzymatic reactions to $y_j$. This process is replicated for each individual pathway and for every sample in $\mathcal{S}$ with four specific constraints (reflecting the rules definitions in PathoLogic [8]):

  1. If only a single enzymatic reaction is attached to $y_j$, we retain that pathway.
  2. If a set of enzymatic reactions is unique to $y_j$, we do not remove those unique reactions.
  3. If $y_j$ is a biosynthesis pathway, we do not remove the last two enzymatic reactions from that pathway.
  4. If $y_j$ is a biodegradation pathway, we do not remove the first two enzymatic reactions from $y_j$.

Because the set of pathways, as defined in MetaCyc database, is unique, distinct, and reflects only a small part of the earth's still unexplored organismal diversity, the *pathway corruption* technique is adopted to create various forms of true pathways that might be encountered in the experimental data due to the errors propagated from the upstream data analysis. Obviously, in creating the synthetic dataset, the above procedure neglects completely the true biological rules; nonetheless, this dataset will provide a separate unbiased measurement on the performance of mlLGPR. In the experiments, we created two synthetic datasets: *Synset-1* that follows Phase 1 and 2 of the generation process while the *Synset-2* adds the Phase 3 on top of the two phases. The number of expected pathways for both data is assumed to follow the Poisson distribution with mean value equal to 500. Note that the corruption is done at the input space, i.e., we assume that $\hat{\mathbf{x}}^{(i)} = \mathbf{x}^{(i)} + \epsilon$, where $\epsilon$ is the amount of corruption. In this paper, we are not concerning to recover the true sample $\mathbf{x}^{(i)}$ and, for brevity, we denote $\hat{\mathbf{x}}^{(i)}$ as $\mathbf{x}^{(i)}$.

### 2 ELA metric

Unfortunately, the performance metrics, such as average F1 score, do not consider the mlLGPR-EN's behavior affected by the noise. Accordingly, we follow the work of Saez et al. [9] and present the *equalized loss of accuracy (ELA)* metric, which quantifies the expected behavior of a model against noise, and defined as:

$$\begin{aligned} &\mathrm{ELA}_\rho = \mathrm{RLA}_\rho + s(M_0) \\ &\text{where } \mathrm{RLA}_\rho = \frac{M_0 - M_\rho}{M_0} \text{ and } s(M_0) = \frac{1 - M_0}{M_0} \end{aligned} \tag{2}$$

The ELA metric embeds both concepts: i)- the robustness of a model, computed by $\mathrm{RLA}_\rho$ at a controlled noise threshold $\rho$ and ii)- the performance of a model without noise, i.e., $s(M_0)$, where 1 represents the base accuracy.

### 3 Experiments

#### 3.1 Pathway Prediction on CAMI data

In this section, we contrast mlLGPR (using elastic net penalty with reaction and pathway evidence features) with PathoLogic generated PGDBs on CAMI low complexity dataset. From Table 2, we observe that mlLGPR achieved an average F1 score of 0.4866 for mlLGPR.

Table 2: **Predictive performance of mlLGPR-EN with AB, RE and PE feature sets on CAMI low complexity data.**

| Metric | mlLGRPR-EN (+AB+RE+PE) |
|---|---|
| Hamming Loss (↓) | 0.0975 |
| Average Precision Score (↑) | 0.3570 |
| Average Recall Score (↑) | 0.7827 |
| Average F1 Score (↑) | 0.4866 |

##### 3.2 Statistical Analyses of Pathway Prediction Algorithms

Inspired by the *Friedman test* [10], in this section, we conduct a systematic approach to compare and rank the pathway prediction algorithms. Let $r_i^j$ denote the rank of the $m$-th of $\mathbf{C}$ algorithms, based on a performance metric, on the $i$-th of $|\mathcal{S}|$ dataset. Also, let $R_m = \frac{1}{|\mathcal{S}|}\sum_i r_i^m$ be the average rank for the $m$-th algorithm under the null-hypothesis that states "all algorithms are equally likely to perform". Then, the Friedman statistic is distributed according to the F-distribution with $\mathbf{C}-1$ and $(\mathbf{C}-1)(|\mathcal{S}|-1)$ degrees of freedom:

$$F_F = \frac{(|\mathcal{S}|-1)\chi_F^2}{|\mathcal{S}|(\mathbf{C}-1)-\chi_F^2} \quad \text{where } \chi_F^2 = \frac{12|\mathcal{S}|}{\mathbf{C}(\mathbf{C}+1)}[\sum_m R_m^2 - \frac{\mathbf{C}(\mathbf{C}+1)^2}{4}] \tag{3}$$

The results of this test are summarized in Table 3. With 7 algorithms and 7 dataset, the critical value of $F_F(6,36)$ for $\tau = 0.05$ significance level is 2.3638, so we reject the null-hypothesis in terms of all metrics because their $F_F$ values are higher than the critical value.

| Metric | $F_F$ | Critical value ($\tau = 0.05$) |
|---|---|---|
| Hamming Loss | 41.4783 | |
| Average Precision | 111.3000 | 2.3638 |
| Average Recall | 57.5250 | |
| Average F1 | 32.1111 | |

Table 3: **Summary of the Friedman statistics $F_F$ for 7 algorithms and 7 datasets.** The critical value $\tau$ is set to 0.05 significance level.

Consequently, we proceed with a *Nemenyi (post-hoc)* [10] test to analyze the relative performance among the pathway prediction algorithms where the mlLGPR-EN is treated as the control algorithm:

$$\text{Critical Difference (CD)} = q_\tau \sqrt{\frac{\mathbf{C}(\mathbf{C}+1)}{6|\mathcal{S}|}} \tag{4}$$

where $q_\tau = 2.949$ at significance level $\tau = 0.05$ and thus CD $= 3.4052$ ($\mathbf{C}=7, |\mathcal{S}|=7$) (see the paper [10]). Henceforth, the performance between mlLGPR-EN and each comparing algorithm is assumed to be significantly different if the corresponding average rank over 7 datasets differs by at least three CD. Fig 2 shows the CD diagrams for four evaluation metrics at 0.05 significance level, where the average rank of each comparing algorithm is marked along the axis. In each sub-figure, the algorithms which are not significantly different are interconnected with a thick line. In summary, among 49 comparisons (7 methods $\times 7$ dataset), all variants of mlLGPR statistically outperforms against all the other methods in terms of Hamming loss and average F1. With regard to average precision, all variants of mlLGPR achieve statistically comparable performances with PathoLogic, however, mlLGPR-EN and mlLGPR-L1 can be compared with BASELINE, Naïve, and MinPath in terms of average recall. These observations indicate the competitive performance of mlLGPR-EN, against the rest of the pathway prediction algorithms, in all of the evaluation metrics.

Figure 2: **Comparison of seven methods against each other with the Nemenyi test using CD diagrams.** Groups of methods that are not significantly different (at $\tau = 0.05$) are connected. (a)- CD diagram for Hamming loss. (b)- CD diagram for average precision score. (c)- CD diagram for average recall score.(d)- CD diagram for average F1 score.

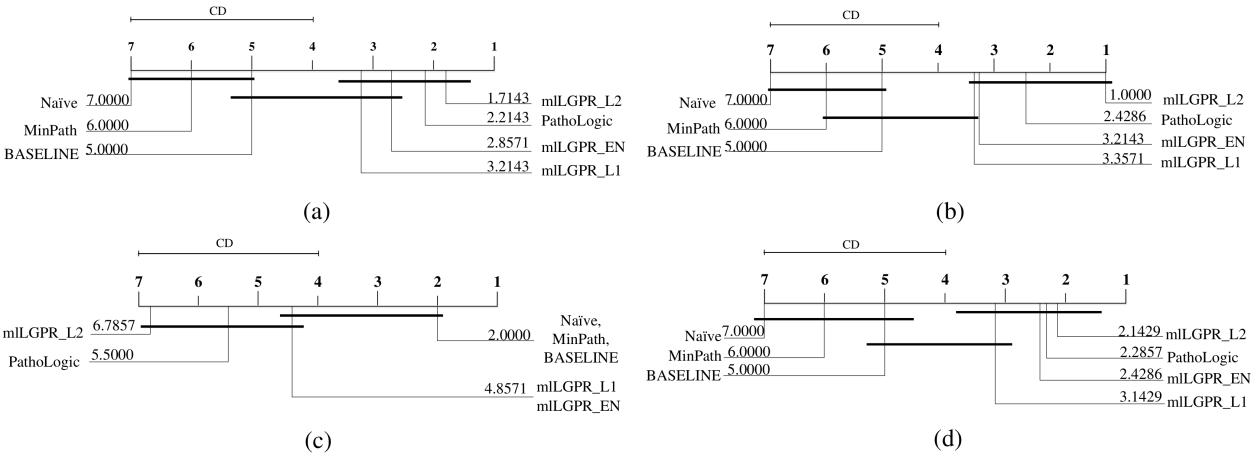
